# A CRISPR/Cas9-induced blunt-end telomere system in *S. pombe* reveals RNase H2-dependent RNA primer removal at the terminal Okazaki fragment of lagging telomeres

**DOI:** 10.64898/2026.07.15.738813

**Authors:** Sai Zou, Tiantian Ye, Lijuan Fu, Jin-Qiu Zhou

## Abstract

Studying the fine-scale dynamics of telomere replication has been hindered by the heterogeneity of native telomeres and the limitations of existing tools. Here, we report a highly efficient and inducible CRISPR/Cas9-mediated system for generating *de novo* telomeres with defined blunt ends in *S. pombe*. This fine setting allows for the precise dissection of post-replicative telomere end structures at near single-nucleotide resolution. Using this system, we show that the replicated leading-strand telomere is blunt-ended, while the lagging-strand counterpart contains a ∼10-nt 3’ overhang resulting from RNA primer removal. By analyzing mutants deficient in ribonucleases, we found that the removal of this terminal RNA primer is specifically dependent on RNase H2, but not RNase H1. This RNase H2-dependent mechanism is essential for defining the mature structure of the lagging-strand telomere with authentic telomeric sequences. Our findings reveal a fundamental asymmetry in telomere end processing after replication and establish RNase H2 as the key enzyme responsible for resolving the terminal RNA primer in lagging-strand telomeres. This mechanism, conserved from budding to fission yeast, underscores the critical and evolutionarily ancient role of RNase H2 in defining eukaryotic telomere architecture.

**GRAPHICAL ABSTRACT:** 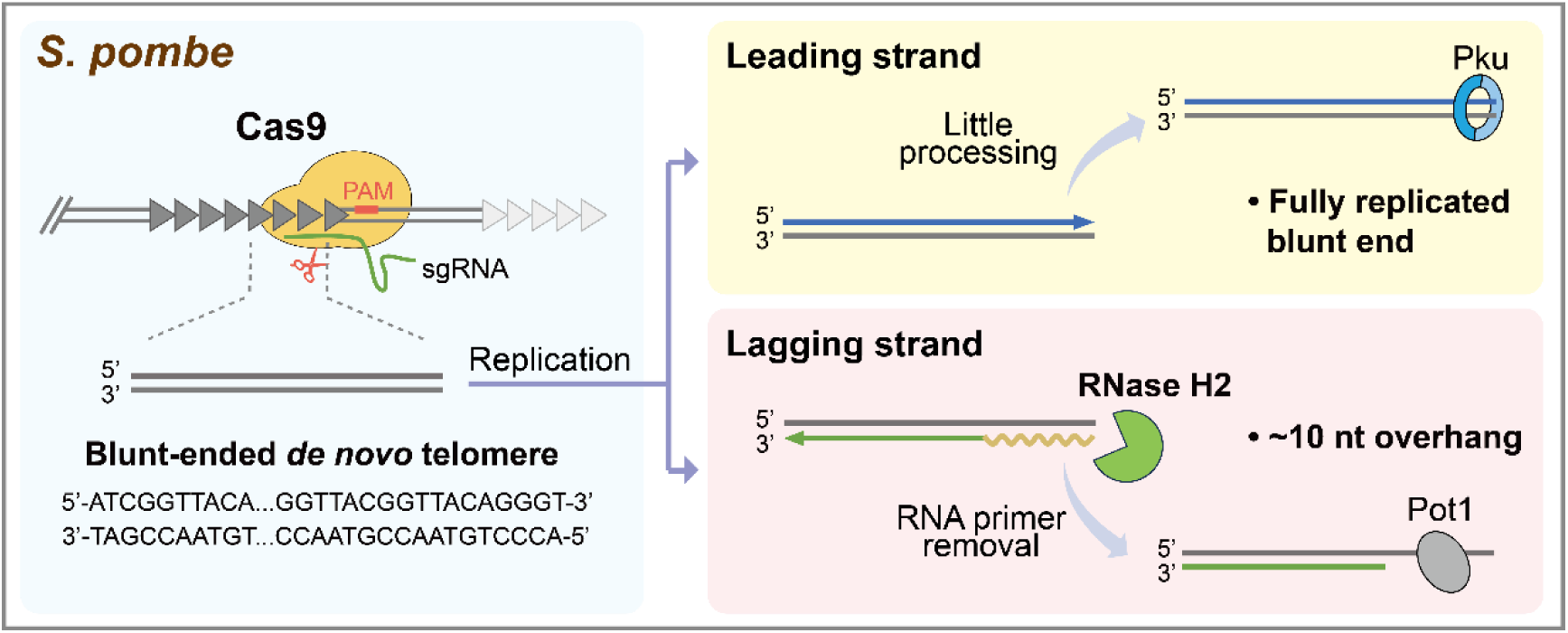

## INTRODUCTION

Telomeres are the physical ends of linear chromosomes in eukaryotic cells and consist of repetitive DNA sequences bound by specific proteins. Their special structure distinguishes chromosome ends from normal DNA double-strand breaks (DSBs) and prevents terminal fusion, repair or degradation, thereby maintaining chromosome integrity and genome stability (1–3).

The telomeric DNA sequence consists of short, guanine-rich nucleotide repeats that vary between species (4). In the fission yeast *Schizosaccharomyces pombe*, the consensus sequence of telomeric DNA is 5’-G_2-6_TTAC(A)-3’/3’-C_2-6_AATG(T)-5’ and it is approximately 300 bp in length (1, 5). The guanine-rich strand, known as the G-strand, usually has a 3’ terminal protrusion (also called a 3’ overhang), which is bound and protected by a specific telomeric single-stranded binding protein (i.e. Pot1) (1, 4). Fission yeast telomeres also contain sub-telomeric regions of approximately 100 kb in length. Part of this region is a 50 kb sub-telomeric homologous (SH) region. The remaining distal part consists of unique sequences that form a highly condensed chromatin structure called the “knob” structure (6).

Telomeres share many structural similarities with DNA double-strand breaks, but the structurally and functionally conserved protein complexes that specifically bind to telomeres (also known as shelterin) maintain the stability of the telomere structure while effectively preventing nucleases from recognizing and degrading the telomere ends (2, 7). In fission yeast, shelterin consists of six proteins: Taz1, Rap1, Poz1, Tpz1, Pot1 and Ccq1. Taz1 and Rap1 form one sub-complex bound to double-stranded DNA, while Tpz1 and Pot1 form another sub-complex bound to single-stranded DNA (8–11). Poz1 bridges these two sub-complexes (11). Ccq1 interacts with Tpz1 to recruit telomerase (11–13). In addition to shelterin, the heterodimeric Pku complex (consisting of Pku70 and Pku80) exerts a protective function at telomeres. Inactivation of the Pku proteins results in shortened telomeres and frequent chromosomal rearrangements in the proximal telomere sequence (14–16). Due to the structural similarities between telomeres and DNA double-strand breaks, many proteins involved in DNA double-strand break repair also function at telomeres (17). For instance, the Mre11-Rad50-Nbs1 complex interacts with Ctp1 to create a single-stranded DNA gap, which initiates the cleavage of the 5’ end of the CA strand by the nuclease Exo1, which in turn produces a 3’ overhang (18–20).

DNA replication relies on DNA polymerases to carry out semi-conservative and semi-continuous replication in a strict 5’ to 3’ direction. The same principle applies to telomeric DNA replication (21–24). The leading strand of telomeric DNA is synthesized continuously in the 5’-3’ direction by DNA polymerase ε (Polε), which uses the C strand as a template. Meanwhile, the lagging strand is synthesized as Okazaki fragments using the G strand as a template (Supplemental Fig. S1). First, Polα-primase synthesizes an 8-12 nucleotide (nt) RNA primer and a ∼20 nt DNA primer, and then Polδ extends the Okazaki fragment synthesis until it displaces the RNA-DNA primer in the downstream Okazaki fragment, forming a flap structure (Supplemental Fig. S1). This structure is subsequently cleaved by the specific nucleases Fen1 (or Dna2). The single-stranded nicks are then sealed by DNA ligase I (Ligase I), completing lagging strand synthesis (25–27). However, when the replication process reaches the very end of the chromosome, there is no upstream Okazaki fragment that displaces the RNA-DNA primer in the very terminal Okazaki fragment. Thus, how the terminal RNA primer is removed remains unclear (27, 28) (Supplemental Fig. S1).

Due to the highly repetitive nature of telomere DNA sequences, the heterogeneity of telomere lengths among different chromosomes and the limited resolution of existing biochemical approaches, it has long been difficult for researchers to monitor the dynamic changes in telomere structure during DNA replication (29–31). Previous studies have shown that the *de novo* telomeres generated by I-SceI endonuclease cleavage can efficiently bind telomere-associated proteins such as Taz1, Pot1 and Ccq1, thereby distinguishing themselves from ordinary DSBs and fulfilling functional roles (32–34). Recently, we used the *de novo* telomere formation system to analyze telomere end replication and protection in the budding yeast *Saccharomyces cerevisiae*, and found that the RNA primer at the terminal Okazaki fragment is removed by RNase H2 (35). However, given the significant evolutionary divergence (approximately 300 to 400 million years) (36) and notable differences in telomere-associated proteins between fission and budding yeast (11, 37–40), it remained an open question whether this RNase H2-dependent mechanism is conserved in *S. pombe*, whose shelterin complex more closely resembles that of mammals. Addressing this question would not only resolve the mechanism of terminal primer removal but also offer insights into the evolutionary constraints shaping telomere replication.

In this study, we developed a highly efficient system for CRISPR/Cas9-mediated chromosome engineering in *S. pombe*. We observed that the conditional induction of CRISPR/Cas9 targets an intrachromosomal telomeric sequence to generate a *de novo* telomere with a blunt end. Additionally, inactivation of RNase H2 results in the retention of the RNA primer in the terminal Okazaki fragment at replicated lagging-strand telomeres, demonstrating that RNase H2 is responsible for the removal of the last RNA primer at *S. pombe* telomeres.

## MATERIAL AND METHODS

### REAGENTS AND TOOLS TABLE

**Table 1.** Reagents and tools used in this study.

| Reagent/Resource | Reference or Source | Identifier or Catalog Number |
| --- | --- | --- |
| **Experimental Models: Fission yeast strains** |  |  |
| *h+ ura4-D18 leu1-32 his3-D1 ade6-M210* | Dr. Li-Lin Du (NIBS) | DY1778 |
| *h- ura4-D18 leu1-32 his3-D1 ade6-M210* | Dr. Li-Lin Du (NIBS) | DY1781 |
| DY1781 *leu1-32::pFA-TetR-PenotetS-Cas9-bsdMX-sgRNA11 gal1-3':ura4+-dnt-sgRNA11 site-hphMX* | This study | DNTFY171 (DF254) |
| DF254 *exo1Δ::natMX* | This study | DNTFY236 |
| DF254 *pku70Δ::kanMX* | This study | DNTFY323 |
| DF254 *pku70Δ::kanMX exo1Δ::natMX* | This study | DNTFY327 |
| DF254 *taz1Δ::natMX* | This study | DNTFY374 |
| DF254 *pku70Δ::kanMX taz1Δ::natMX* | This study | DNTFY378 |
| DF254 *rnh201Δ::natMX* | This study | DNTFY175 |
| DF254 *rnh1Δ::kanMX* | This study | DNTFY258 |
| DF254 *rnh1Δ::kanMX rnh201Δ::natMX* | This study | DNTFY260 |
| DY1778 *leu1-32::pFA-TetR-PenotetS-Cas9-bsdMX-sgRNA15* | This study | DNTFY240 |
| DY1778 *leu1-32::pFA-TetR-PenotetS-Cas9-bsdMX-sgRNA15 gal1-3':ura4+-dnt-sgRNA15 site-kanMX* | This study | DNTFY297 (DA260) |
| DA260 *rnh201Δ::natMX* | This study | DNTFY252 |
| DA260 *rnh1Δ::hphMX* | This study | DNTFY355 |
| DA260 *rnh1Δ::hphMX rnh201Δ::natMX* | This study | DNTFY359 |
| **Recombinant DNA** |  |  |
| Plasmid: pFA-TetR-PenotetS-Cas9-bsdMX-sgRNA11 | This study | pDNT55 |
| Plasmid: pRS314-ura4+-dnt-sgRNA11-hphMX | This study | pDNT69 |
| Plasmid: pRS314-ura4+-dnt-sgRNA15-hphMX | This study | pDNT97 |
| Plasmid: pFA-TetR-PenotetS-Cas9-bsdMX-sgRNA15 | This study | pDNT107 |
| **Oligonucleotides and other sequence-based reagents** |  |  |
| Oligonucleotides used in the study see details in Supplemental Tables S3-5 | Azenta | - |
| **Chemicals, Enzymes and other reagents** |  |  |
| Propidium Iodide | Sigma | P4170 |
| Hydroxyurea | Sigma | H8627 |
| Proteinase K | Sigma | P6556 |
| Hygromycin B | Sigma | V900372 |
| L-Glutamic acid | Sigma | G5889 |
| Single-stranded template DNA | Sigma | D9156 |
| Nylon membrane | Thermo Scientific | AM10104 |
| RNase A | Thermo Scientific | EN0531 |
| EcoRV | Thermo Scientific | FD0304 |
| SuperSignal™ West Atto | Thermo Scientific | A38556 |
| Chemiluminescent Nucleic Acid Detection Module Kit | Thermo Scientific | 89880 |
| Exo I | NEB | M0293L |
| RNase H | NEB | M0297L |
| Electrophoresis mode | Cytiva | SE260 |
| Blasticidin S | Abcam | ab141452 |
| Nourseothricin sulfate | Solarbio | N9210 |
| G418 sulfate | BBi | A600958-0005 |
| Anhydrotetracycline | APExBio | C4291 |
| DNA extraction reagent | Psaitong | PS1013 |
| Zymolyase | MP Biomedicals | 08320921 |
| Acryl/Bis 30% Solution (29:1) | Sangon Biotech | B546017 |
| Ligation high Ver.2 | TOYOBO | LGK-201 |
| In-Fusion HD Cloning Kit | Takara | 639649 |
| **Software** |  |  |
| sgRNA design software | zlab | https://www.benchling.com/crispr |
| Fiji/ImageJ | Fiji/ImageJ | https://imagej.net/ij/ |
| R, 4.4.1 | The R project | https://www.r-project.org/ |
| FlowJo version 10.2 | BD Biosciences | https://docs.flowjo.com/flowjo/getting-acquainted/10-2-release-notes/ |

## METHODS AND PROTOCOLS

### Strain construction

All yeast strains (see details in Table 1) of *de novo* telomere system are derivatives of the *Schizosaccharomyces pombe* strain: *ura4-D18 leu1-32 his3-D1 ade6-M210*, which was obtained from Dr. Li-Lin Du (National Institute of Biological Sciences (NIBS), China). Cas9 expression cassette (TetR-NLS-Cas9-NLS*-bsdMX-*sgRNA) was inserted to the *leu1-32* locus. *De novo* telomere cassette (*ura4^+^-dnt-hphMX/kanMX*) was inserted to the 2-kb region 3’ of the *gal1^+^* gene 3’ untranslated region (UTR) (32). Gene knock-in was performed by a PCR-amplified DNA product-mediated homologous recombination followed by Lithium acetate transformation, and positive clones were verified by PCR, and the resulting strain hereafter served as the WT control strain. Gene deletion mutants were generated by PCR fragments mediated gene knockout.

### *De novo* telomere induction and cell cycle monitoring

All strains were first inoculated in hygromycin- or G418-containing selection medium to monitor potential leaky expression, and were then cultured to log phase.

For *de novo* telomere formation assay, after being cultured to log phase in hygromycin/G418-containing medium, cells were collected and washed three times in EMM-N medium (with low supplements) to remove residual nitrogen sources, and then resuspended in EMM-N medium. After 16 hours (h) of synchronization, ahTET was added to induce Cas9 expression. Cells were harvested at the indicated time points.

For cell replication experiment, cells were collected and washed with EMM-N medium and resuspended in EMM-N medium. After 24 h of nitrogen starvation to achieve G0 arrest, ahTET was added to induce Cas9 expression for 8 h during this period. Subsequently, cells were collected and washed twice with sterile water to remove residual ahTET, then resuspended in YES medium to release cells into the cell cycle. Cells were harvested at different time points. Cell cycle progression was confirmed by FACS analysis. All cell culture experiments were conducted at 32 ℃. See Table 1 for thereagents used.

### Flow cytometry (FACS)

Cells were first washed once with sterile ddH_2_O, then thoroughly mixed with 70% ethanol and incubated for 1 h at room temperature or overnight at 4 ℃. Fixed cells were washed twice with 50 mM sodium citrate buffer (pH 7.2) and resuspended in the same buffer containing RNase A (final concentration of 200 µg/ml), followed by incubation at 37 ℃ for 2-3 h. Proteinase K (final concentration of 200 µg/ml) was added, followed by incubation at 50 ℃ for 1 h. Cells were washed and diluted to an appropriate concentration, followed by sonication and stained with propidium iodide (PI, 10 µg/ml) for 1 h at room temperature. FACS analysis was then performed (41). See Table 1 for the reagents used.

### Genomic DNA extraction

Cells collected from each time point were stored at −80 ℃, and their genomic DNA was extracted in parallel using a mild protocol (42). Cells were first digested with zymolyase at a final concentration of 1.5 mg/ml in sorbitol buffer, then resuspended in lysis buffer, followed by phenol/chloroform extraction and ethanol precipitation. Finally, genomic DNA was dissolved in an appropriate volume of ddH_2_O. See Table 1 for the reagents used.

### Southern blot analysis

The dissolved DNA was further purified by incubation with RNase A. Additional RNase H treatment was performed at 37 ℃ for 2 h in the RNA primer detection assay. Extracted DNA was digested with EcoRV at 37℃ for 2 h, then Exo I was added, followed by incubation at 37 ℃ for 1 h to remove the 3’ overhang.

For native Southern blotting (35), the final digested DNA was separated on native 8% PAGE and transferred to a nylon membrane (Thermo Fisher). Then crosslinked by UV and incubated with denature buffer (0.5 M NaOH, 0.5% SDS) at 45 ℃ for 30 min, washed three times with ddH_2_O, 5 minutes each time, and finally hybridized with CA probe (5’-TGT AAC CCC TGT AAC CCC TGT AAC CCC-3’-biotin) at 50 ℃ overnight. Detection of the signal was performed according to the manufacturer’s instructions (Chemiluminescent Nucleic Acid Detection Module, Thermo Fisher). Finally, the membrane was exposed to the CCD camera.

For denaturing Southern blotting (35), an equal volume of 2×denature loading buffer (95% formamide, 0.025% bromophenol blue, 0.025% xylene cyanol FF, 0.5 mM EDTA) was added to the final digestion product and boiled at 100 ℃ for 10 min, then immediately chilled into ice for 5 min. Single-stranded DNA was then separated on a denaturing urea (8 M)-6% PAGE, transferred to membrane, crosslinked by UV and hybridized with the specific CA or TG probe as indicated. Signal detection was performed as described for native PAGE. The sequences of the 3’-biotin-labelled probes are as follows: CA probe, 5’-TGT AAC CCC TGT AAC CCC TGT AAC CCC-3’-biotin; TG probe, 5’-GGG GTT ACA GGG GTT ACA GGG GTT ACA-3’-biotin. Sequences of the size markers for denaturing and native PAGE are listed in Supplemental Tables S3-S5, respectively. All results were repeated three times independently. See Table 1 for the reagents used.

### Double-stranded and single-stranded markers preparation

For denaturing PAGE markers, the oligonucleotides used in this study were synthesized by the Azenta company. The sequences of the oligonucleotides are listed in Supplemental Tables S3-S5, which closely matched E-TRF sequences.

For native PAGE markers, dsDNA fragments were generated by annealing the complementary oligonucleotides used for denaturing PAGE. The annealing reaction was heated to 95 ℃ for 10 min, then cooled from 95 ℃ to 65 ℃ at a rate of 0.01 ℃ per second, and held at 65 ℃ for 5 min. The annealing products were separated on an agarose gel and extracted from the gel.

### Quantification and statistical analysis

Quantification in Figure 1D of the telomere signals was performed by the ImageJ. Data are shown as mean ± SEM and n represents the total number of independent biological replicates. For each lane, the signal intensities of the target bands and control bands were normalized to their respective probe-hybridizing lengths, and the normalized target signal was divided by the normalized control signal to obtain a normalized ratio. The bar plots were created using Microsoft Excel 2021.

**Figure 1.**
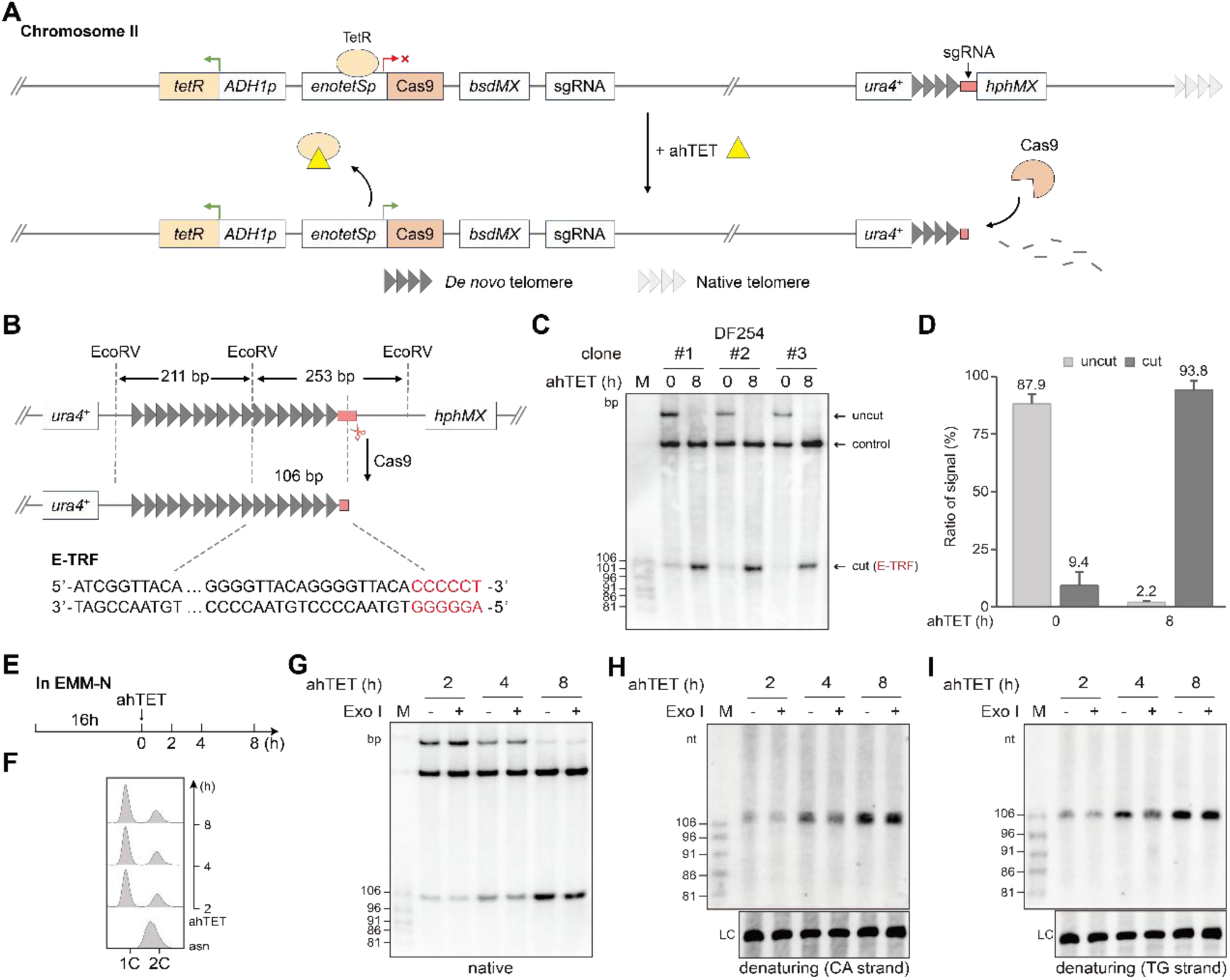
*De novo* telomere generated by CRISPR/Cas9 possesses a stable blunt end. **A**, Schematic illustration of the CRISPR/Cas9-mediated *de novo* telomere induction system. Constitutive expression of TetR inhibits *enotetS* promoter activity. The addition of ahTET induces Cas9 expression and cuts the sgRNA target sequence (pink rectangle), exposing the *de novo* telomere (filled triangles), which ends with a 6 bp non-telomeric sequence (shown in red in panel B). **B**, A zoomed-in view of the *de novo* telomere cassette containing the *ura4^+^*gene, followed by 254 bp of telomere repeats and the hygromycin resistance marker (*hphMX*). EcoRV recognition sites are present both within and flanking the *de novo* telomere sequence. Cas9 cleavage results in a 106 bp E-TRF with a blunt end. The sequence of the E-TRF is displayed at the bottom, with the Cas9 cleavage site highlighted in red (non-telomeric sequence). **C**, Southern blot detection of CRISPR/Cas9-mediated *de novo* telomere formation in WT cells, with 253 bp “uncut”, 211 bp “control” and 106 bp “cut” (E-TRF) bands. **D**, Quantitative analysis of the signal ratios of uncut and cut bands before and after ahTET induction. Signal intensities were normalized to the corresponding control band after normalization for the length of the probe-hybridizing region. Data are presented as mean ± SEM, *n* = 3/group. **E**, Flow chart of *de novo* telomere formation (see text for a detailed description). Cells are collected at the indicated time points for FACS and Southern blot analysis. **F**, FACS analysis of the DNA content of nitrogen-starvation synchronized WT cells. asn: asynchronous. **G**, Southern blot detection of the 3’ overhang of the *de novo* telomere (E-TRF) induced in WT cells. **H** and **I**, Denaturing Southern blot detection of the CA-strand (H) and TG-strand (I), which represent the same membrane hybridized with TG and CA probe, respectively. LC: loading control.

Quantification in Figures 3E and 4I of the telomere signals was performed by the ImageJ software. The net signal intensities of the Exo I-sensitive and Exo I-resistant daughter telomere bands were quantified after background subtraction and normalized to the length of the corresponding probe-hybridizing region. The percentage of each daughter telomere was determined by dividing the normalized signal intensity of the corresponding band by the total normalized signal intensity of the Exo I-sensitive and Exo I-resistant daughter telomere bands. Data are presented as mean ± SEM and n represents the total number of independent biological replicates. Statistical significance was determined by two-way ANOVA followed by Tukey’s multiple comparisons test. Significance levels were indicated as: *p < 0.05, **p < 0.01, ***p < 0.001, ns, not significant. The bar plots were created using Microsoft Excel 2021.

## RESULTS

### A CRISPR/Cas9-induced *de novo* telomere system generates a stable blunt end

Endonucleases such as I-SceI and HO have been widely used to create DSBs for exposing *de novo* telomeres (32, 43). However, these systems suffer from low cleavage efficiency and generate complex end structures. For instance, I-SceI cleavage produces DSBs containing approximately 9 bp of non-telomeric sequences with a 4-nt 3’ overhang, raising concerns about whether such ends can be processed as authentic telomeres. To overcome these limitations and establish a more robust and precise tool for fission yeast telomere biology, we developed a novel inducible CRISPR/Cas9 system. This system generates *de novo* telomeres with defined blunt ends, mirroring a potential native telomere state and enabling high-resolution downstream analyses.

We combined a previously reported Cas9 expression system with the Tet-On system (44–46) to construct a system for rapid induction of *de novo* telomere formation in *S. pombe* (Fig. 1A). The *enotetSp*/Cas9-*rrk1*/sgRNA expression cassette and a *tetR* gene (encoding TetR repressor) cassette (under the control of an *adh1* promoter) were integrated into the *leu1-32* locus on chromosome II (Fig. 1A). Additionally, a 254 bp telomere sequence flanked by *ura4*^+^ and the sgRNA recognition sequence-*hphMX* gene was inserted near the native right telomere on chromosome II (Fig. 1A). Presumably, the constitutively expressed TetR repressor binds to the *enotetS* promoter, blocking Cas9 expression. When ahTET is added into the medium, it competitively binds to TetR, which derepresses the *enotetS* promoter, and induces the expression of the Cas9 (flanked by NLS sequences at both ends). The expressed Cas9/sgRNA complex recognizes the target sequence proximal to the 254 bp telomere sequence and generates a double-stranded break, exposing a *de novo* telomere with a blunt end. It is worth noting that the optimized *de novo* telomeric sequence was 254 bp long; such a long telomere could circumvent telomerase extension (47–49), a conclusion also supported by our own data (Fig. 1C, Supplemental Fig. S2). The *de novo* telomeric sequence consists of regular 5’-GGGGTTACA-3’ repeats, with one EcoRV site in the middle (Supplemental Table S1). Two additional EcoRV restriction sites were designed at both ends of the telomeric sequence to facilitate the detection of the EcoRV-digested terminal restriction fragment (E-TRF) by Southern blotting (Fig. 1B). When Cas9 is not induced, EcoRV digestion yields a control band of 211 bp and an uncut band of 253 bp (Fig. 1B). Upon Cas9 induction, a cut band (E-TRF) of 106 bp appears (Fig. 1B).

After testing multiple sgRNA sequences and evaluating their cleavage efficiencies (Supplemental Fig. S2, Supplemental Table S2), we selected the DF254 strain for further experiments (Fig. 1C, Supplemental Fig. S3). Induction of Cas9 with ahTET resulted in efficient cutting of the target sequence, exposing the *de novo* telomere which displayed the anticipated 106 bp E-TRF (Fig. 1C). Quantitative analysis showed that after 8 h of ahTET induction, nearly all uncut bands were cleaved and converted into the E-TRF band (Fig. 1D). These results demonstrate that we have established a novel inducible CRISPR/Cas9 system for rapid and highly efficient genome editing in *S. pombe*.

To determine the terminal structure of *de novo* telomeres generated by Cas9, we arrested cells in the G0 phase by nitrogen starvation for 16 h (46) (Fig. 1E and F), then induced Cas9 expression by adding ahTET and harvested the cells at various time points. The *de novo* telomere structure was analyzed by Southern blotting. Results from both native and denaturing gels showed that the E-TRF maintained a constant length of 106 bp/nt in the absence or presence of Exo I digestion (Fig. 1G-I). These results suggest that *de novo* telomeres induced by Cas9 have blunt ends.

### The Pku complex and endogenous Exo1 regulate end resection at *de novo* telomeres

The *S. pombe* Pku complex binds to telomere ends and plays a role in maintaining telomere stability (14–16). To investigate whether the Pku complex inhibits the resection of the CA strand of *de novo* telomeres, we constructed a DF254 *pku70*Δ strain and analyzed the structure of *de novo* telomere ends in cells arrested in the G0 phase (Fig. 2A). When the E-TRFs of the *de novo* telomeres were examined in a native gel by Southern blotting, they were 106 bp in length within 2-8 h of induction (Fig. 2B). However, a subset of E-TRFs were sensitive to Exo I digestion, displaying diffuse bands ranging from approximately 91 to 106 bp (Fig. 2B), suggesting that short-range resection had occurred at the *de novo* telomeres (Supplemental Fig. S4). Denaturing Southern blotting further revealed that the CA strand exhibited a prominent 106-nt band at the 8 h induction time point, both before and after Exo I treatment, along with several additional faint bands of approximately 100, 91, and 81 nt in length. This pattern was not significantly altered by Exo I digestion (Fig. 2C). In contrast, prior to Exo I treatment, the TG strand was predominantly 106 nt in length; after Exo I digestion, it displayed a similar band pattern to that of the CA strand (Fig. 2D). These results collectively indicate that in the absence of Pku complex protection, the 5’ end of the CA strand of the *de novo* telomere becomes more susceptible to nuclease resection.

**Figure 2.**
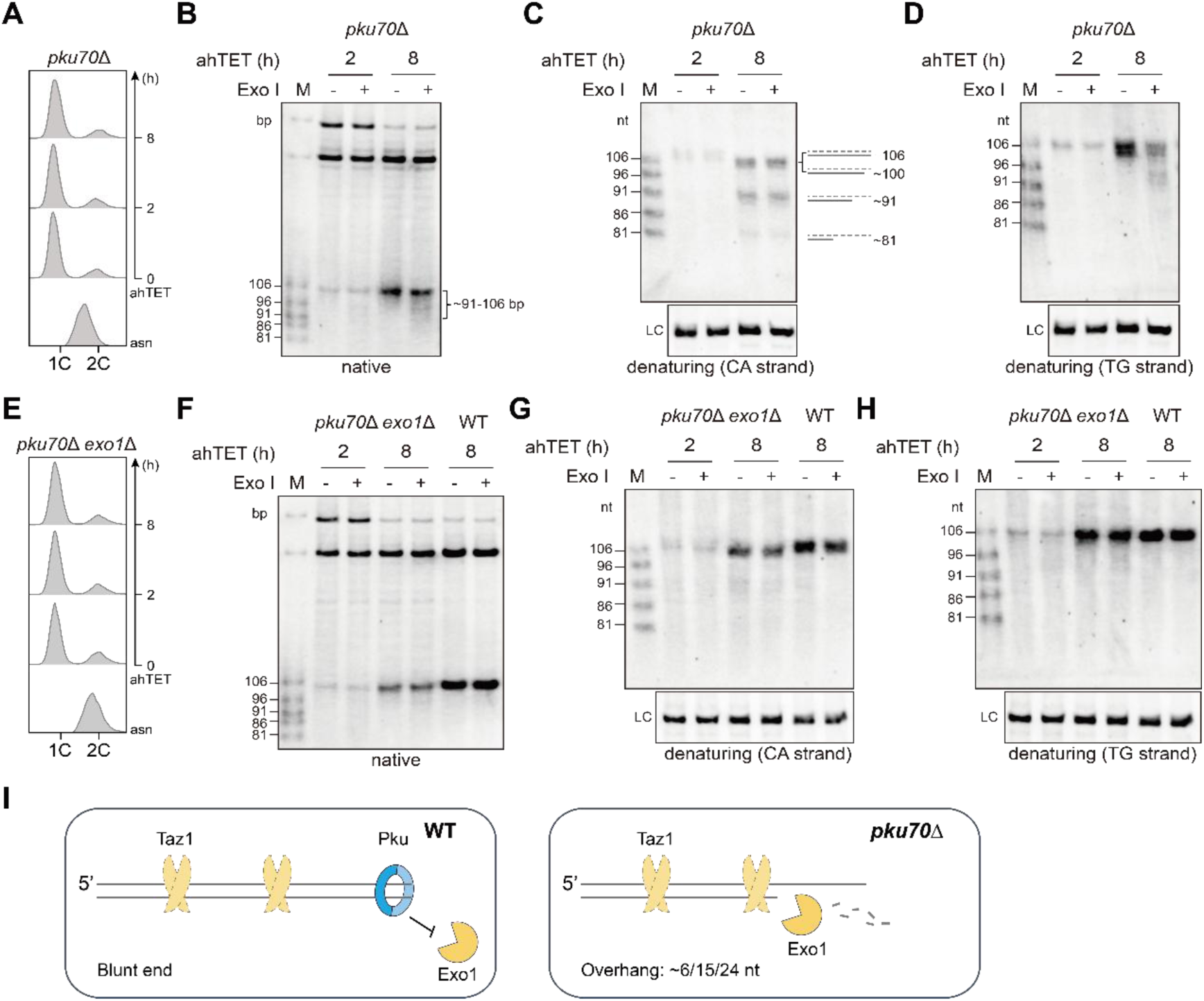
*De novo* telomere is protected by Pku complex, preventing from endogenous Exo1-mediated resection. **A** and **E**, FACS analysis of the DNA content of nitrogen-starvation synchronized *pku70*Δ (A) and *pku70*Δ *exo1*Δ (E) cells. **B** and **F**, Southern blot detection of the 3’ overhang of the *de novo* telomere induced in *pku70*Δ (B) and *pku70*Δ *exo1*Δ (F) cells. **C** and **D**, Denaturing Southern blot detection of the CA strand (C) and TG strand (D) in *pku70*Δ cells, which represent the same membrane hybridized with TG and CA probe, respectively. LC: loading control. **G** and **H**, Denaturing Southern blot detection of the CA strand (G) and TG strand (H) in *pku70*Δ *exo1*Δ cells, which represent the same membrane hybridized with TG and CA probe, respectively. LC: loading control. The WT lanes shown in F-H correspond to the same Southern blots as those shown in B-D, respectively. They are presented separately for clarity. **I**, Model for *de novo* telomere end protection and processing in WT and *pku70*Δ cells.

We next deleted the *exo1* gene, which encodes the 5’-3’ exonuclease Exo1 that is involved in telomere 5’ resection (50, 51), in both the DF254 and DF254 *pku70*Δ strains, and examined the *de novo* telomere structure using native and denaturing Southern blot assays. Native Southern blot results showed that in *exo1*Δ cells (a negative control), the *de novo* telomere end structure was as expected (Supplemental Fig. S5A, C, D and E), identical to that in WT cells (Fig. 1G-I). Importantly, in the *pku70*Δ *exo1*Δ cells, the 5’ resection of the CA strand observed in *pku70*Δ cells (Fig. 2B) was no longer detected (Fig. 2F). Consistently, denaturing gel results further showed that both the CA and TG strands in the *pku70*Δ *exo1*Δ strain remain 106-nt in length, indistinguishable from those in WT cells, indicating little or no resection (Fig. 2G and H). These results collectively support the model that, in the absence of the Pku complex protection, the CA strand of the exposed *de novo* telomere is susceptible to Exo1. In other words, the Pku complex protects *de novo* telomere ends from Exo1-mediated resection, thereby avoiding the generation of long 3’ overhang structures (14, 50–52) (Fig. 2I). Notably, compared to WT strain, faint signals below the 106 nt band were detected in the *pku70*Δ *exo1*Δ strain, suggesting that additional nucleases may also contribute to CA strand resection of the *de novo* telomere when Pku is absent.

Why is the resection of the CA strand in *pku70*Δ cells restricted to specific lengths, rather than occurring as a continuous size range? We reasoned that the telomeric double-stranded DNA-binding protein Taz1 (9, 53) acts as a physical barrier to protect the ends from resection (Supplemental Fig. S4). The ladder pattern of the CA strand in Figure 2C correlates well with the sporadic presence of Taz1 binding sites in the *de novo* telomere sequence. For example, a terminal Taz1 binding site lies just eight nucleotides from the end of the CA strand (Supplemental Fig. S4). We constructed *taz1*Δ and *pku70*Δ *taz1*Δ mutants and examined *de novo* telomere structures. Unexpectedly, the *de novo* telomere signals in the *taz1*Δ mutant displayed a similar pattern to those in the WT cells (Supplemental Fig. S5 A, C, D and E). Additionally, resection of the *de novo* telomere was barely detectable in the *pku70*Δ *taz1*Δ cells compared with wild-type cells (Supplemental Fig. S5G-I). Furthermore, faint bands of intermediate size below cut bands were detected in *taz1*Δ strain (Fig. S5D), but their origin could not be determined. They did not appear to be generated by resection, as no products of the same size were detected on the complementary TG strand (Fig. S5E). Because Cas9 cleavage efficiency is markedly reduced in *taz1*Δ cells under our experimental conditions (53), these data do not allow us to directly determine whether Taz1 itself is responsible for the observed resection pattern. Additional experiments will be required to test the role of Taz1 in this process.

### Replication generates asymmetric telomere ends: a blunt leading strand and an overhanging lagging strand

We next examined the structures of daughter telomeres derived from the *de novo* telomere after one round of replication. The experimental protocol was as follows. The cells were cultured in EMM-N medium for 16 h to synchronize them in the G0 phase. Cas9 expression was then induced by adding ahTET for 8 h to generate *de novo* telomeres. The cells were then transferred to YES medium to allow progression through cell cycle until completion of one round of cell cycle (Fig. 3A). To facilitate analysis of replicated daughter telomere structures, we prepared a positive control marker, designated “OM” by mixing two DNA fragments in equal amounts: a 106 bp blunt-ended DNA that is insensitive to Exo I; the other is a 106 bp DNA with a 10-nt 3’ overhang that is sensitive to Exo I (Fig. 3B). The sequences of both fragments closely match that of the E-TRF.

**Figure 3.**
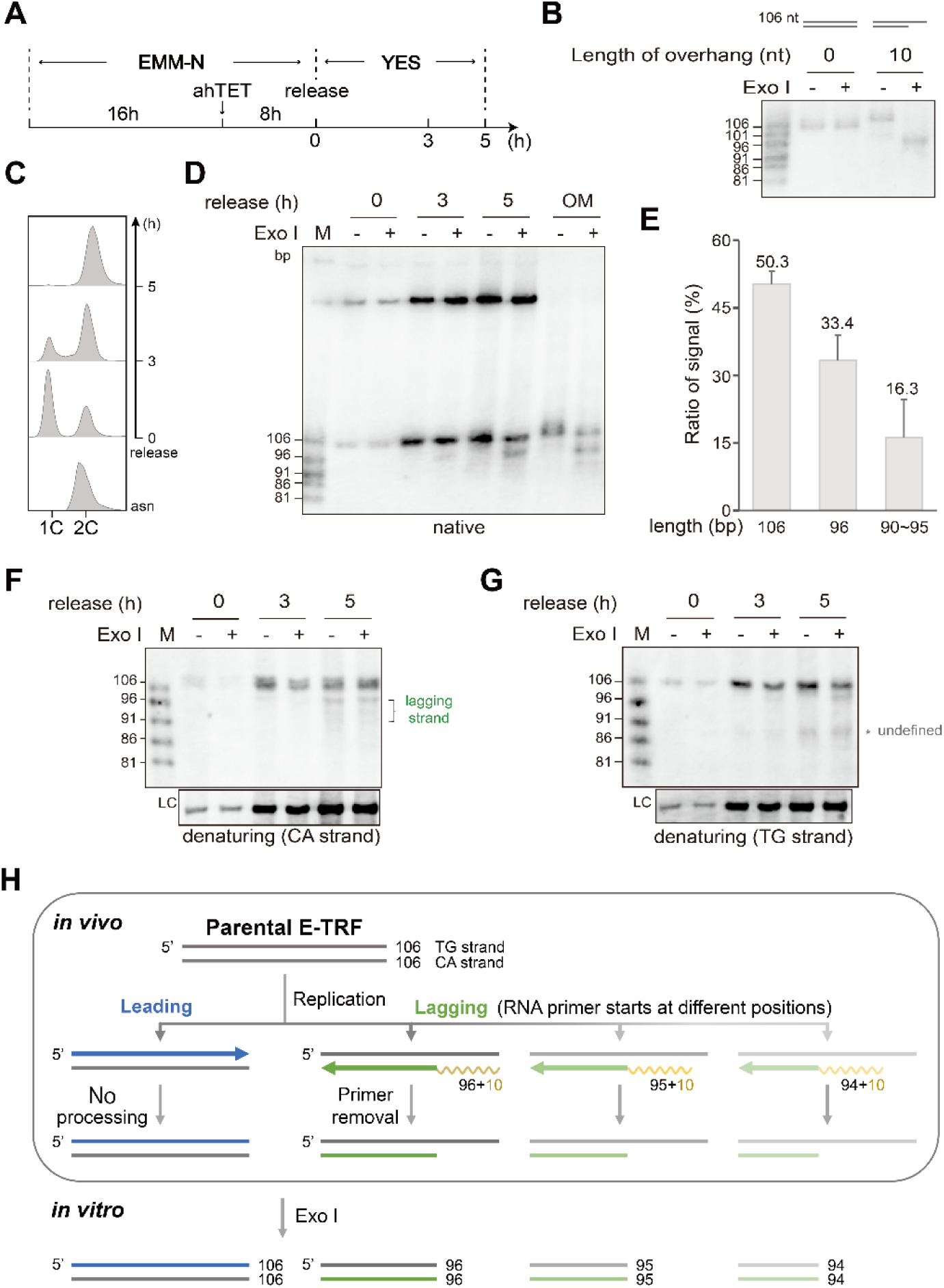
Approximately Half of the daughter telomeres are blunt-ended after DNA replication. **A**, Flow chart of *de novo* telomere replication within one cell cycle (see text for a detailed description). Cells are collected at the indicated time points for FACS and Southern blot analysis. **B**, Southern blot analysis of the 3’ overhang of oligonucleotide markers: a 106 bp blunt-ended DNA and a 106 bp DNA with a 10-nt overhang, as labeled. **C**, FACS analysis of DNA content in WT cells before and after release into the cell cycle. asn: asynchronous. **D**, Southern blot analysis of the 3’ overhangs at both telomere ends during one round of DNA replication in WT cells. OM: oligonucleotide marker, Grey arrow: overlapped daughter telomeres, Blue arrow: blunt-ended leading telomere, Red square bracket: Exo I-trimmed lagging telomere. **E**, Quantification of daughter telomeres with different end structures (E-TRFs in the lane of release 5 h, +Exo I). The percentages of daughter telomeres with different structures were calculated from the background-subtracted signal intensities, normalized to the length of the corresponding probe-hybridizing regions. Data are presented as mean ± SEM, *n* = 3/group. **F** and **G**, Denaturing Southern blot detection of the CA-strand (F) and TG-strand (G) after replication in WT cells, which represent the same membrane hybridized with TG and CA probe, respectively. LC: loading control. **H**, Schematic illustration of the different structures at the leading and lagging daughter telomeres that are replicated from the parental *de novo* telomere: the leading telomere has a blunt end, whereas the lagging telomere has a 3’ overhang after the removal of the last RNA primer. The structures of the lagging strand vary slightly due to different positions of the RNA primer initiation at the terminus.

The cells were harvested at specific time points and their DNA content was monitored by FACS analysis. The “2C” peak appeared at 3 h and became dominant at 5 h, indicating the completion of DNA replication (Fig. 3C). The lengths and structures of the E-TRFs were then examined using Southern blotting. As shown in Figure 3D, the E-TRF of the parental telomere measured 106 bp and was insensitive to Exo I treatment prior to replication (Fig. 3D, 0 h), confirming blunt end as mentioned above in Figure 1. After replication, the E-TRFs of the daughter telomeres were 106 bp in length prior to Exo I digestion (Fig. 3D, 5 h, -Exo I). Strikingly, upon Exo I digestion, the population of replicated E-TRFs resolved into two distinct species: one maintaining the parental 106 bp size (blunt-ended), and the other represented the resection products, which were mainly present as an approximately 96-bp band, accompanied by a minor smear signal ranging from 90 to 95 bp (Fig. 3D, 5 h, +Exo I). This size difference is consistent with the *in vitro* trimming of a ∼10 nucleotide 3’ overhang from a subset of the telomeres. Quantitative analysis revealed the 106-bp signal accounted for 50.3% of the total, while the resection products collectively accounted for 49.7% (the 96-bp band comprising 33.4% and the 90-95 bp smear comprising 16.3%). Given the inherent asymmetry of DNA replication, we reasoned that the two species likely correspond to the leading- and lagging-strand replication products (Fig. 3H). Thus, the E-TRF of lagging telomere contains a double strand region and a 10-15-nt 3’ overhang (Fig. 3H). This 10-15 nt overhang likely corresponds to the region previously occupied by the RNA primer.

The lengths of both the replicated TG and CA strands were further examined using denaturing Southern blotting with CA and TG probes, respectively. Prior to Exo I treatment, the TG-strand signal was predominantly 106 nt at each time point (Fig. 3G). After Exo I treatment, approximately half of the TG-strand signal converted to 96 nt or shorter after replication (Fig. 3G, 5 h, +Exo I). We speculate that the bands labeled “undefined” (Fig. 3G) result from a small fraction of *de novo* telomeres that escaped protection and were subsequently resected. The CA-strand signal measured 106 and 96 nt after replication (Fig. 3F, 5 h), regardless of Exo I digestion. Notably, faint but diffuse CA-strand signals below the 96-nt band were visible. We speculate that these signals originate from differences in RNA primer initiation during terminal Okazaki fragment synthesis, as RNA primer synthesis can begin from either the first thymidine or the cytosines proximal to the 3’ end due to the nucleotide preference of primases (Fig. 3H) (54, 55). These results are consistent with the native Southern blot data, and suggest the following: (1) the replicated leading strand telomere undergoes little (or no) end processing and maintains a stable blunt-end structure. (2) The replicated lagging strand telomere contains a 3’ overhang of approximately 10-15 nt. (3) The RNA primer at the terminal Okazaki fragment can be synthesized from the end of the TG strand, and is removed after replication. (4) The length of the RNA primer is approximately 10 nt. Taken together, this finding reveals a fundamental asymmetry in telomere end processing after replication in *S. pombe*.

### The RNA primer in the terminal Okazaki fragment on the lagging strand telomere is removed after completion of DNA replication

During DNA replication, DNA Polα-primase synthesizes an RNA-DNA primer to initiate Okazaki fragment synthesis on the lagging strand (26, 56). In the telomeric region, the RNA primer at the end of the Okazaki fragment is thought to be removed, exposing the 3’ overhang and allowing Pot1 to bind and protect the telomere end (Supplemental Fig. S1) (11, 57). To address whether RNA primers are present in the very termini of replicated lagging strands, we prepared genomic DNA in the presence or absence of RNase H (Supplemental Fig. S6), an enzyme that digests the RNA moiety in DNA-RNA hybrids *in vitro* (58, 59). Native Southern blot results showed that the hybridization signals of replicated daughter telomeres exhibited the same pattern regardless of RNase H treatment (Fig. 4A and B). The lagging strand signal slightly increased from 45.3% to 51.8% upon RNase H treatment, but the difference was not statistically significant (Fig. 4I). These data suggest that the RNA primers in the terminal Okazaki fragments are efficiently removed during or after telomere replication.

**Figure 4.**
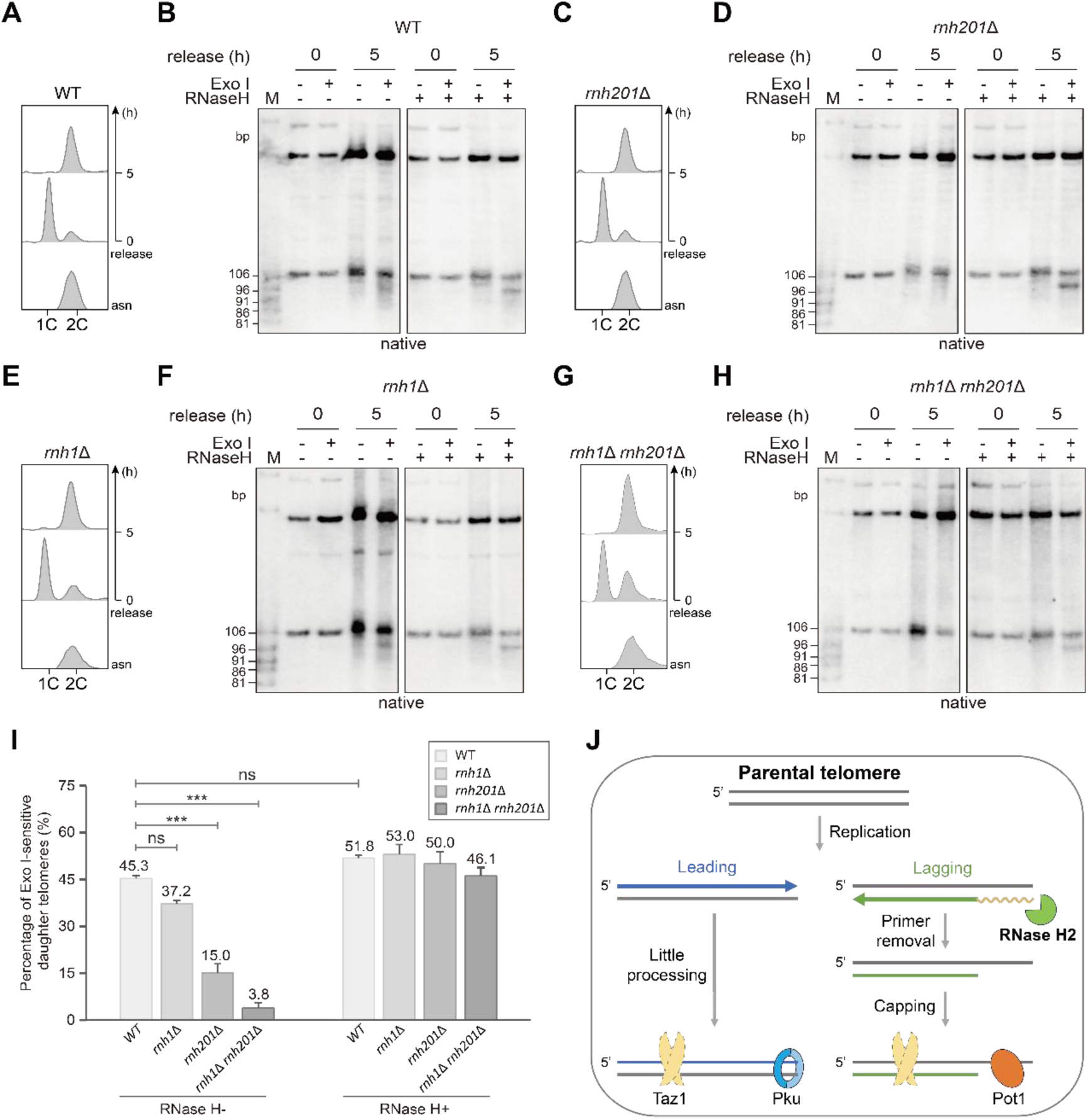
The last RNA primer is retained on lagging-strand telomere in *rnh201*Δ cells. **A**, **C**, **E** and **G**, FACS analysis of DNA content in WT (A), *rnh201*Δ (C), *rnh1*Δ (E) and *rnh1*Δ *rnh201*Δ (G) cells. asn: asynchronous. **B**, **D**, **F** and **H**, Southern blot analysis of the 3’ overhang at both the leading- and lagging-strand telomere ends during one round of DNA replication in WT (B), *rnh201*Δ (D), *rnh1*Δ (F) and *rnh1*Δ *rnh201*Δ (H) cells. Green arrow: Exo I-trimmed lagging telomere. Genomic DNA was pretreated with or without RNase H *in vitro*, as labeled. **I**, Quantification of the percentage of Exo I-sensitive daughter telomeres in WT, *rnh1*Δ, *rnh201*Δ, and *rnh1*Δ *rnh201*Δ cells when pretreated with or without RNase H *in vitro* (E-TRFs in the lane of release 5 h, +Exo I). The percentage of Exo I-sensitive daughter telomeres was calculated from the background-subtracted signal intensities, normalized to the length of the corresponding probe-hybridizing regions. Data are presented as mean ± SEM, *n* = 3/group. Statistical significance was determined by two-way ANOVA followed by Tukey’s multiple comparisons test. Significance levels are indicated as: *p < 0.05, **p < 0.01, ***p < 0.001, ns, not significant. **J**, A model for telomere replication in fission yeast. A blunt-ended parental telomere produces two daughter telomeres: the replicated leading strand telomere undergoes minimal processing, and its blunt end is capped by Pku, whereas the replicated lagging strand telomere has a Pot1-protected overhang following the removal of the terminal RNA primer.

### RNase H2 is specifically required for the removal of the terminal RNA primer

In the internal region of a chromosome, the RNA primer of an Okazaki fragment is typically displaced by an adjacent Okazaki fragment, resulting in the formation of a flap structure. This structure is subsequently cleaved or degraded by endonucleases, such as Fen1 (and/or Dna2) (27, 60–62) (Supplemental Fig. S1). However, the RNA primer at the very end of the lagging strand telomere cannot be removed by the same mechanism as an internal RNA primer, because there is no upstream Okazaki fragment to displace it (Supplemental Fig. S1). As endoribonucleases RNase H1 and RNase H2 can digest the RNA moiety in a DNA-RNA hybrid and play a role in the removal of RNA primers in lagging strand DNA maturation (63–65), it has long been assumed that the primers at the terminal ends of lagging telomeres are removed by RNase H1 and/or RNase H2. However, due to the rarity and short length of these terminal RNA primers, they have been extremely difficult to detect, and this issue has remained experimentally unverified.

To genetically identify the nuclease responsible for terminal primer removal in *S. pombe*, we analyzed strains deficient in RNase H1 (*rnh1*Δ) or the catalytic subunit of RNase H2 (*rnh201*Δ). In the absence of RNase H1 and without *in vitro* RNase H treatment, the replicated lagging-strand telomeres remained sensitive to Exo I with a slight reduction in signal intensity compared to wild-type cells (Fig. 4E, F and I). In contrast, loss of RNase H2 resulted in a marked loss of Exo I sensitivity, indicating a failure to generate the 3’ overhang (Fig. 4C and D left, -RNase H). These results suggest that the RNA primer at the terminus of the lagging telomere is retained in the *rnh201*Δ mutant, but not in the *rnh1*Δ mutant. The definitive proof came from an *in vitro* RNase H rescue experiment: pretreatment of *rnh201*Δ samples with RNase H restored Exo I sensitivity to approximately half of the telomeres (Fig. 4D right, +RNase H), demonstrating that the obstruction to overhang formation was indeed the retained RNA primer. Thus, in *S. pombe*, RNase H2 is both necessary and sufficient for the removal of the terminal RNA primer. Further combinatorial deletion of *rnh1* and *rnh201* resulted in nearly complete retention of the RNA primer on the replicated lagging strand telomere (Fig. 4G and H). We therefore conclude that RNase H2 is the primary enzyme responsible for removing the terminal RNA primers from lagging strand telomere, while RNase H1 may play a minor auxiliary role (Fig. 4J).

### A *de novo* telomere with an authentic telomeric sequence produces a blunt-ended daughter leading strand and a 3’ overhanging daughter lagging strand

The *de novo* telomere generated by Cas9 in the DF254 strain ends with the 6 bp sequence 5’-CCCCCT-3’/3’-GGGGGA-5’ (Supplemental Fig. S3), which differs greatly from the telomeric consensus sequence 5’-G_2-6_TTAC(A)- 3’/3’-C_2-6_AATG(T)-5’. It could be argued that the replication of this non-telomeric sequence and the associated end processing might not recapitulate the events that occur on native telomeres. To unequivocally demonstrate that our findings are not an artifact of the initial non-canonical terminal sequence but rather reflect a general mechanism for authentic telomeres, we engineered a new strain (DA260) where Cas9 cleavage exposes a native *S. pombe* telomeric sequence 5’-GGTTACAGGGT-3’/3’-CCAATGTCCCA-5’ (TG strand/CA strand) (Fig. 5A), matching the consensus native telomeric sequence 5’-G_2-6_TTAC(A)- 3’/3’-C_2-6_AATG(T)-5’. We then examined both the length and the structure of the *de novo* telomeres during one round of replication (Fig. 5B and C). The E-TRFs with a native telomeric sequence at the end measured 106 bp, and were insensitive to Exo I treatment prior to replication, indicating a blunt end (Fig. 5C, 0 h). After replication, smeared signals below 106 bp were detected after Exo I digestion (Fig. 5C, 5 h, +Exo I). These results were similar to those obtained from the DF254 strain (Fig. 4B). Further *in vitro* RNase H treatment of the E-TRF samples slightly enhanced the lagging strand band signals from 41.8% to 45.0% (Fig. 5C, Supplemental Fig. S7). The replication of ends with native telomeric sequences produced asymmetric daughter telomere structures, confirming that the generation of a blunt-ended leading strand and an overhanging lagging strand is a fundamental feature of telomere replication in fission yeast.

**Figure 5.**
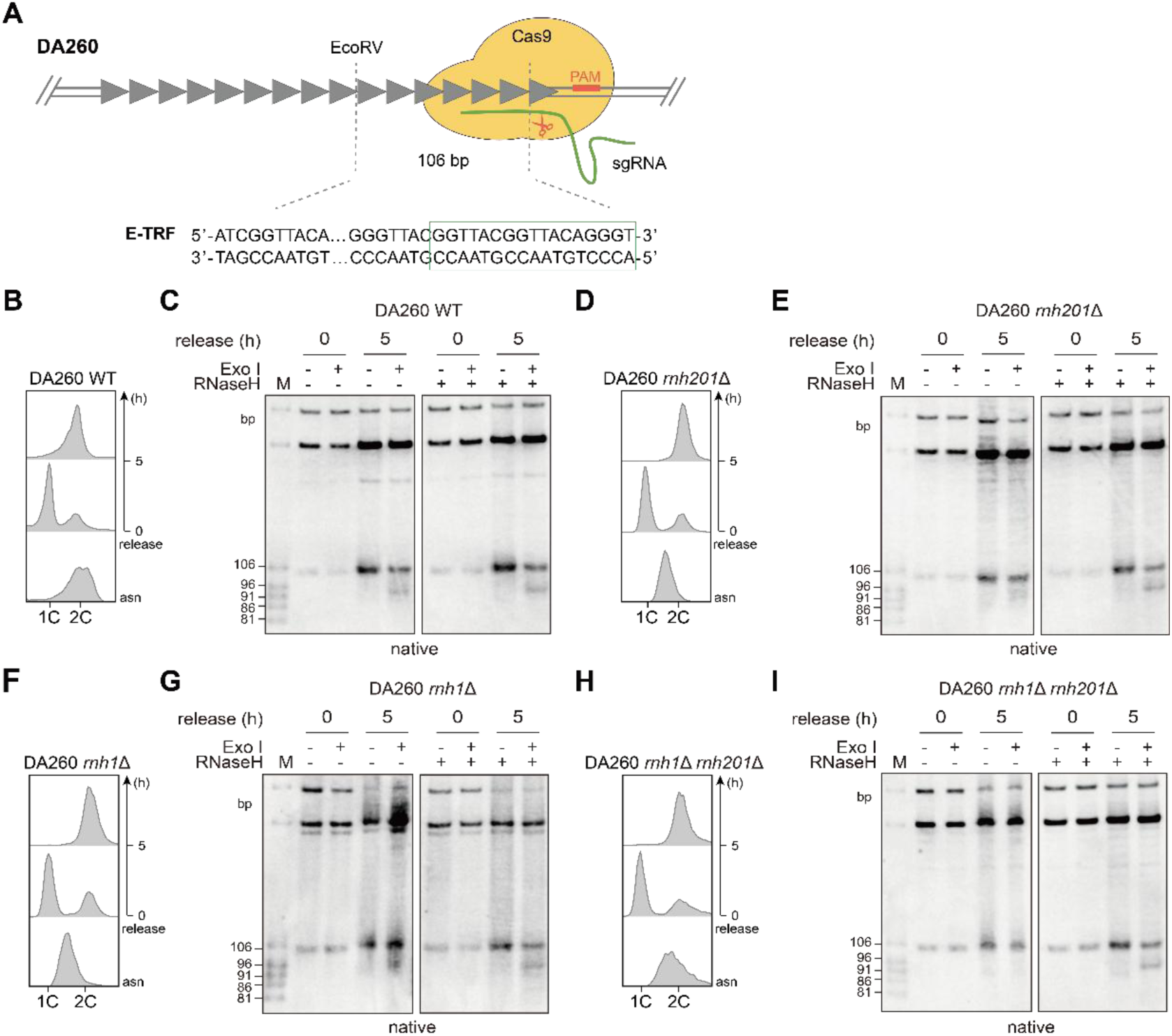
The replication of the *de novo* telomere with an authentic telomeric tip produces a daughter leading telomere with a blunt-ended and a daughter lagging telomere with a 3’ overhang. **A,** Schematic illustration of the CRISPR/Cas9-mediated formation of *de novo* telomere with an authentic telomeric sequence (displayed at the bottom). The sgRNA pairs with a typical telomere sequence immediately upstream of its PAM sequence (boxed in red at the bottom). Cas9 cleavage exposes the new telomere ending in the sequence 5’-GGTTACGGTTACAGGGT-3’/3’- CCAATGCCAATGTCCCA-5’ (in green rectangle). **B**, **D**, **F** and **H**, FACS analysis of the DNA content in DA260, DA260-*rnh201*Δ, DA260-*rnh1*Δ and DA260-*rnh1*Δ *rnh201*Δ cells before and after release into the cell cycle. asn: asynchronous. **C**, **E**, **G** and **H**, Southern blot analysis of the 3’ overhang at both the leading and lagging telomere ends during one round of DNA replication in DA260 (C), DA260- *rnh201*Δ (E), DA260-*rnh1*Δ (G) and DA260-*rnh1*Δ *rnh201*Δ (H) cells. Green arrow: Exo I-trimmed lagging telomere. Genomic DNA was pretreated with or without RNase H, as labeled.

We also deleted *rnh1* and/or *rnh201* in the DA260 strain and examined the structures of the leading and lagging daughter telomeres replicated from Cas9-induced *de novo* telomeres (Fig. 5D, F and H). The results from the *rnh1*Δ strain were nearly identical to those observed in WT DA260 strain (Fig. 5G), In both the DF254 *rnh201*Δ and *rnh1*Δ *rnh201*Δ strains, all of the E-TRFs of the replicated *de novo* telomeres were 106 bp regardless of Exo I treatment (Fig. 5E and I, 0 h), indicating that both the leading and lagging strand telomeres are blunt-ended. When the E-TRFs were pretreated with RNase H and then digested by Exo I *in vitro*, a lagging strand signal appeared in addition to a 106 bp band (Fig. 5E and I, 5 h, +Exo I, +RNase H), indicating that the RNA primer is retained on the lagging telomere, i.e., the lagging strand ends with a blunt DNA-RNA primer hybrid, when RNase H2 is absent. These results consistently support the conclusion that RNase H2 is responsible for removing the terminal RNA primers from lagging strand telomeres (Fig. 4J).

## DISCUSSION

Telomeres play a critical role in maintaining genome stability (1, 7). Studying the mechanism of telomere replication and protection is highly significant for understanding fundamental biological issues such as genome stability, cellular senescence, and cell fate regulation. However, the highly repetitive nature of telomere sequences, the heterogeneity of telomere lengths within and between cells, and the limitations of existing telomere length analysis methods have made it difficult for researchers to precisely observe the fine structure of telomeres, particularly during the complex end processing that occurs during DNA replication at telomeres (66–68). Our current study, for the first time presents a significant methodological advance for studying telomere biology in *S. pombe*—a highly efficient and inducible CRISPR/Cas9 system for generating *de novo* telomeres with defined blunt ends. This tool overcomes the limitations of previous endonuclease-based systems (such as I-SceI or HO) (32, 43) and provides unprecedented resolution for dissecting the fine-scale dynamics of telomere replication and end-processing. In addition, compared with previously reported CRISPR/Cas9 systems, such as those using the *nmt41* promoter to regulate Cas9 expression, which has a complicated procedure and exists a 14-15 h long lag (44, 45), or the *adh1* promoter-driven Cas9 expression (69), which causes cytotoxicity in fission yeast, our current optimized CRISPR/Cas9 system provides a powerful and generally applicable tool for the fission yeast community (Supplemental Fig. S8), enabling high-resolution analysis of telomere dynamics and other time-sensitive chromosomal processes.

Applying this system, we have delineated the molecular events that follow the replication of a telomere in fission yeast, revealing both conserved principles and unique protective mechanisms. A central finding of our work is the inherent structural asymmetry of replicated telomeres: the leading strand telomere is maintained as a stable blunt end, while the lagging strand telomere bears a ∼10 nt 3’ overhang (Figs. 3-5). This challenges the long-held assumption that leading strand telomeres undergo extensive processing and fill-in synthesis to achieve a uniform overhanging structure (2, 70). Crucially, we demonstrate that this blunt-end structure is not an artifact of our experimental system, as it is faithfully produced even when the *de novo* telomere terminates with authentic native telomeric sequences (Fig. 5). The stability of the leading strand blunt end is likely ensured by the protective capping function of the Pku complex, which we show acts as a critical barrier to Exo1-mediated resection (Fig. 2).

Perhaps the most mechanistically insightful finding is the identification of RNase H2 as the primary enzyme responsible for removing the terminal RNA primer on the lagging strand telomere. This role of RNase H2 is specifically conserved in *S. pombe*, as RNase H1 plays little detectable role (35) (Figs. 4 and 5). The elegant genetic evidence from our mutant analyses: the RNA primer is retained in *rnh201*Δ cells and can be subsequently removed *in vitro* by RNase H (Figs. 4 and 5), provides compelling support for this conclusion. Together with our parallel findings in *S. cerevisiae* (35), this establishes RNase H2-mediated RNA primer removal as a fundamental and evolutionarily conserved cause to the “end replication problem” across a vast evolutionary distance, separating budding and fission yeast by over 300 million years. (Fig. 4J).

The conservation of this mechanism in *S. pombe* carries particular significance. Unlike *S. cerevisiae*, the fission yeast telomeric shelterin complex shares greater homology with that of mammals (2, 4, 38). Therefore, our findings may offer a more direct and relevant model for understanding telomere replication in higher eukaryotes. The existence of blunt-ended leading strand telomeres has been reported in plants and mammals (71, 72), but the mechanisms governing their formation and protection have remained unclear. Our work in *S. pombe* provides a clear mechanistic framework: full replication yields a blunt end, which is then stabilized by capping complexes like Ku (Pku in yeast), obviating the need for further processing. It will be of great interest to determine whether RNase H2 performs the same indispensable role in resolving terminal Okazaki fragments in mammalian cells. It is worth noting that the length of RNA primers in fission yeast has long been extrapolated from *in vitro* studies (73, 74). Our present study zooms in the length of the primer at the lagging strand telomere of fission yeast and provides *in vivo* evidence to confirm the widely accepted claim that RNA primers are approximately 10 nucleotides in length (Figs. 3 and 4).

In summary, we propose a revised model for telomere replication (“the asymmetrical end replication model”) where the end replication problem is predominantly driven by the inevitable removal of the RNA primer from the lagging strand, with the leading strand being fully replicated and protected (Fig. 4J). The CRISPR/Cas9-based *de novo* telomere system we developed provides a powerful genetic tool for the fission yeast community, enabling future high-resolution studies not only of telomere biology but also of other chromosomal dynamics. Finally, given the links between RNase H2 dysfunction and human autoimmune disorders and genome instability (75, 76), our findings raise the intriguing possibility that defects in telomere replication may contribute to these disease pathologies.

## ACKNOWLEDGMENTS

We thank lab members for helpful discussion. This work was support by grants from the National Key Research and Development Program of China (No. 2023YFA0913400) and the Shanghai Academy of Natural Sciences (SANS).

## AUTHOR CONTRIBUTION

J.-Q.Z. conceived the study, analyzed the data and wrote the manuscript. S.Z. performed all of the experiments and data analysis. T.Y. established the methodological framework. L.F. involved in reagents preparation and discussed the work. J.-Q.Z. supervised the project.

## DATA AVAILABILITY

This study includes no data deposited in external repositories.

