## Supplemental Figure S1-S8, Table S1-S5 for "A CRISPR/Cas9-induced blunt-end telomere system in *S. pombe* reveals RNase H2-dependent RNA primer removal at the terminal Okazaki fragment of lagging telomeres"

#### Supplemental information

Supplemental figures S1-S8

Supplemental tables S1-S5

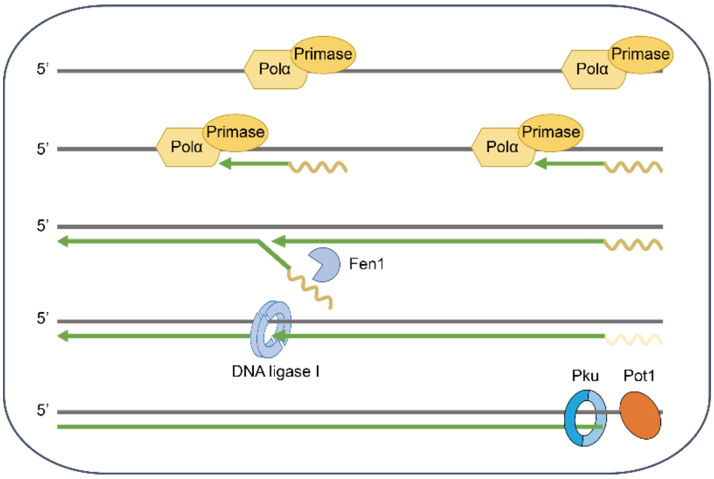


**Supplemental Figure S1. Schematic illustration of lagging strand DNA replication and the terminal processing of a telomere on the lagging strand.**

The lagging strand of DNA is synthesized in discontinuous segments known as Okazaki fragments. Each fragment is initiated by the DNA Polα-primase complex and subsequently joined together by DNA ligase I to form a continuous strand of DNA (see text for a detailed description). In the telomere region, the RNA primer of the final Okazaki fragment is removed, exposing a 3' overhang. This overhang is bound and protected by Pot1, while the junction of the double- and single-stranded DNA is bound and protected by Pku.

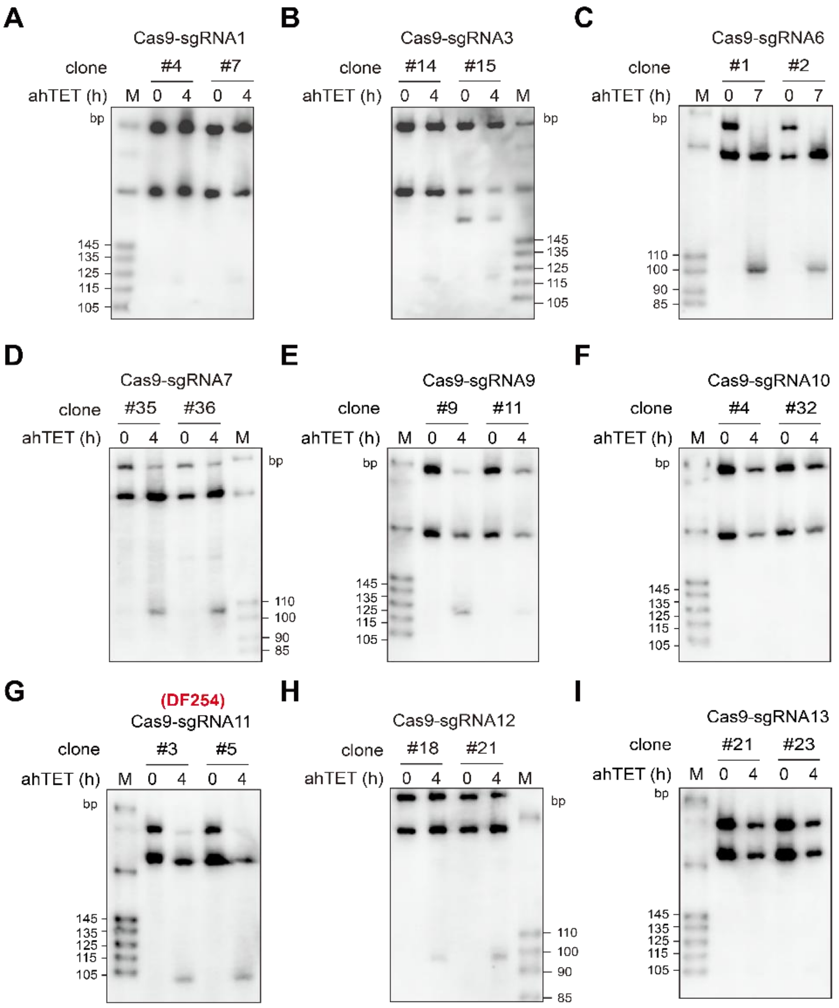


**Supplemental Figure S2. Validation of *de novo* telomere formation mediated by CRISPR/Cas9 with different sgRNA target sequences**

Southern blot detection of *de novo* telomere formation in the strains expressing different sgRNAs (labeled on top of each panel). The *de novo* telomeres were induced in YES medium, and the cutting efficiency was roughly judged by the appearance and intensity of the E-ERF signals. We finally chose strain DF254, which expresses sgRNA11, to perform the subsequent experiments. The sgRNA target sequences are listed in Supplemental Table S2. *De novo* telomere could be induced in the strains showed in **A**, **B**, **C**, **D**, **E**, **G** and **H**, but not in the strains showed in **F** and **I**.

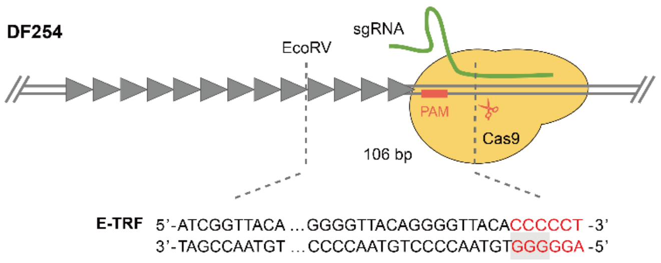


**Supplemental Figure S3. Schematic illustration of the end structure of CRISPR/Cas9-mediated *de novo* telomere in DF254 strain**

The PAM sequence (red rectangle) is adjacent to *de novo* telomere, while the sgRNA pairs with the adjacent non-telomeric region. Cas9 cleavage exposes the telomere, leaving a 6 bp non-telomeric sequence at the very tip. The E-TRF sequence is displayed at the bottom, with PAM sequence boxed in gray; sequence in red represents non-telomeric sequence of Cas9 cleavage.

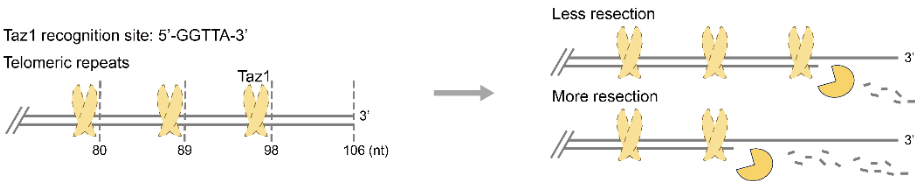


**Supplemental Figure S4. Schematic illustration of potential Taz1 binding at the terminus of *de novo* telomere sequence**

When *pku70* is absent, Exo1-mediated short-range resection takes place, but may be blocked by Taz1.

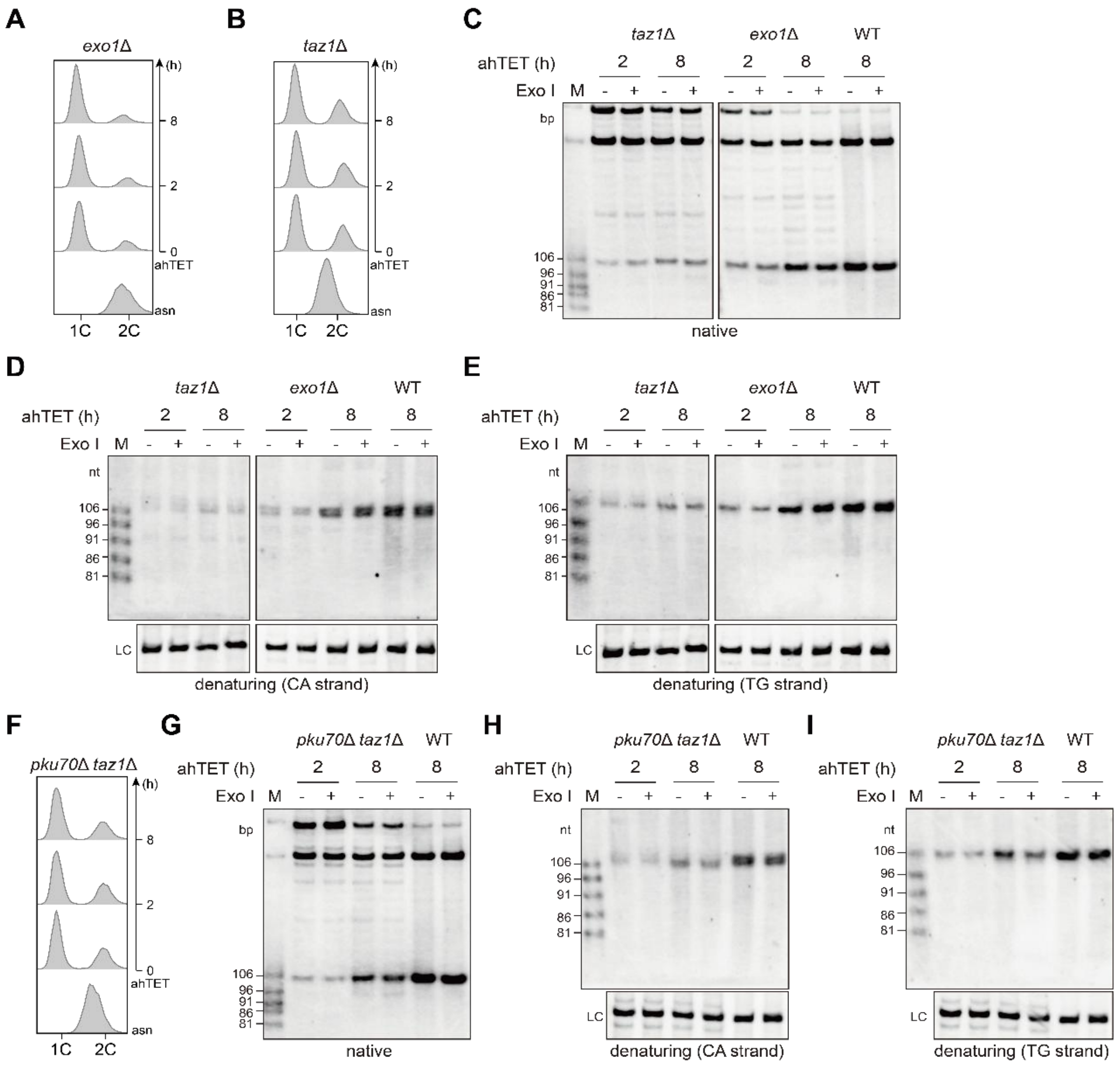


**Supplemental Figure S5. Examination of the length and structure of the *de novo* telomere in *taz1*Δ, *exo1*Δ and *pku70*Δ *taz1*Δ cells**

**A**, **B**, and **F**, FACS analysis of the DNA content of nitrogen-starvation synchronized *exo1*Δ (A), *taz1*Δ (B) and *pku70*Δ *taz1*Δ (F) cells. asn: asynchronous. **C** and **G**, Southern blot detection of the 3' overhang of the *de novo* telomere (E-TRF) induced in *exo1*Δ (C), *taz1*Δ (C) and *pku70*Δ *taz1*Δ (G) cells. **D** and **E**, Denaturing Southern blot detection of the CA-strand (D) and TG-strand (E) in *exo1*Δ and *taz1*Δ cells, which represent the same membrane hybridized with TG and CA probe, respectively. LC: loading control. **H** and **I**, Denaturing Southern blot detection of the CA-strand (H) and TG-strand (I) in *pku70*Δ *taz1*Δ cells, which represent the same membrane hybridized with TG and CA probe, respectively. LC: loading control.

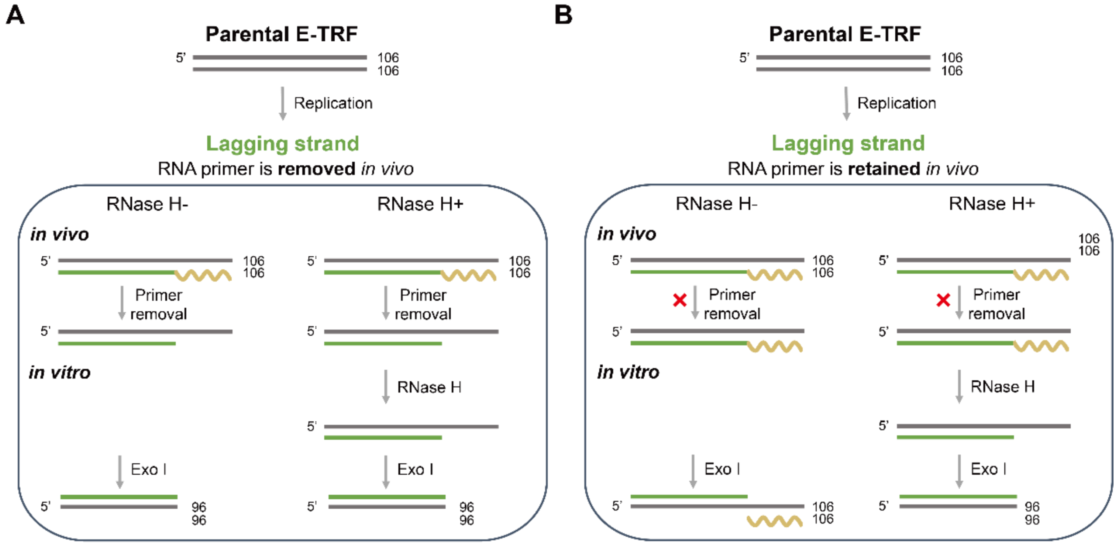


**Supplemental Figure S6. Schematic illustration of the last RNA primer detection**

**A**, *In vivo* RNA primer (yellow squiggly line) removal after replication leaves an overhang of ~10 nt on lagging strand. The genomic DNA shows similar sensitivity to *in vitro* Exo I digestion in the absence or presence of RNase H. **B**, RNA primer is retained after replication, the lagging strand ends with a blunt DNA-RNA primer hybrid. The genomic DNA is insensitive to *in vitro* Exo I digestion when not treated with RNase H (left panel), but becomes sensitive to *in vitro* Exo I digestion because of a 3' overhang generated by RNase H (right panel).

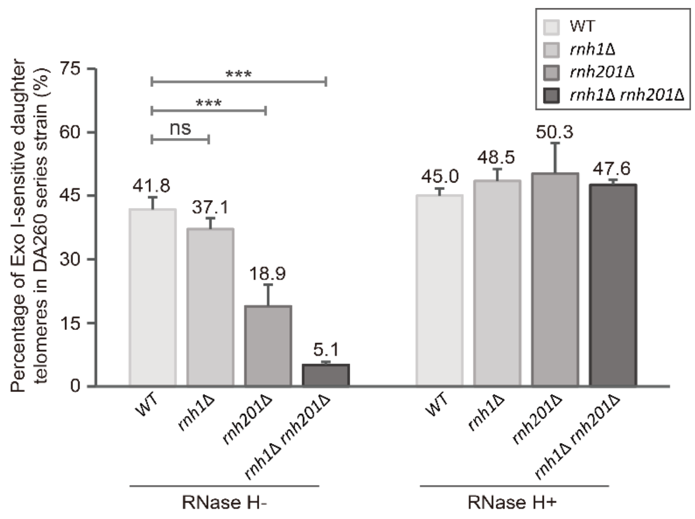


**Supplemental Figure S7. Quantitative analysis of the ratio of lagging strand in DA260 series strain**

Quantification of the percentage of Exo I-sensitive daughter telomeres in DA260-WT, -*rnh1*Δ, -*rnh201*Δ, -*rnh1*Δ *rnh201*Δ cells when pretreated with or without RNase H *in vitro* (E-TRFs in the lane of release 5 h, +Exo I). The percentage of Exo I-sensitive daughter telomeres was calculated from the background-subtracted signal intensities, normalized to the length of the corresponding probe-hybridizing regions. Data are presented as mean ± SEM, n = 3/group. Statistical significance was determined by two-way ANOVA followed by Tukey's multiple comparisons test. Significance levels are indicated as: *$p < 0.05$, **$p < 0.01$, ***$p < 0.001$, ns, not significant.

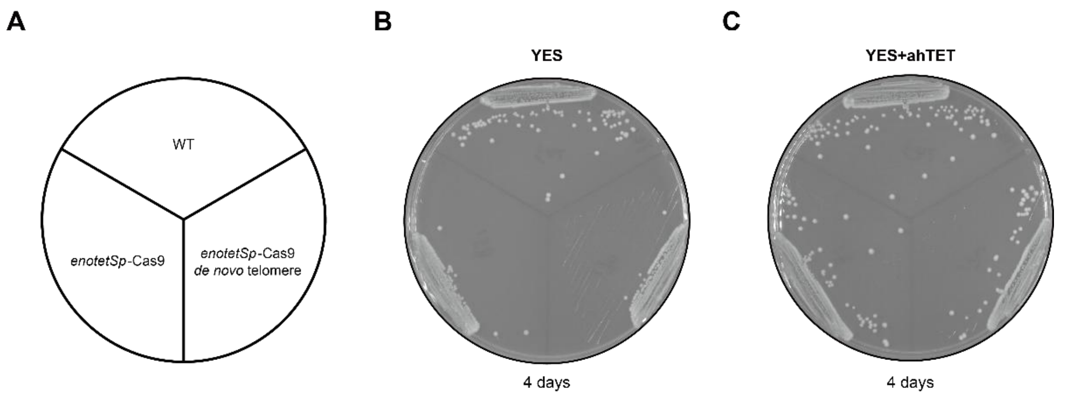


**Supplemental Figure S8. Growth analysis of the strains that expresses Cas9**

**A**, Upper: wild type strain (Cas9-null); Lower left: strain with Cas9 expression under the control of *enotetS* promoter; Lower right: strain with Cas9 expression under the control of *enotetS* promoter, capable of cleaving and exposing *de novo* telomere. **B**, Three strains grown on YES plate after 4-day incubation. **C**, Three strains grown on YES-ahTET plate after 4-day induction of Cas9.

### A CRISPR/Cas9-induced blunt-end telomere system in *S. pombe* reveals RNase H2-dependent RNA primer removal at the terminal Okazaki fragment of lagging telomeres

Sai Zou[1], Tiantian Ye[1], Lijuan Fu[1,2] and Jin-Qiu Zhou[1,2*]

[1]State key Laboratory of RNA innovation-Science and Technology, CAS Center for Excellence in Molecular Cell Science, Shanghai Institute of Biochemistry and Cell Biology, Chinese Academy of Sciences; University of Chinese Academy of Sciences, Shanghai 200031, China

[2]School of Life Science and Technology, ShanghaiTech University, 201210 Shanghai, China

#### Supplemental information-tables

Supplemental tables S1-5

**Supplemental Table S1: Sequences of designed *de novo* telomere after cleaved by Cas9**

| | Length (bp) | Sequence (5'-3') |
|---|---|---|
| DF254 (dsDNA) | 254 | TTACAGGGGTTACAGGGGTTACAGGGGTTACAGGGGTTACAGGGG<br>TTACAGGGGTTACAGGGGTTACAGGGGTTACAGGGGTTACAGGGG<br>TTACAGGGGTTACAGGGGTTACAGGGGTTACAGGGGTTACAGGGG<br>TTACAGGGGTTACAGGGATATCGGTTACAGGGGTTACAGGGGTTAC<br>AGGGGTTACAGGGGTTACAGGGGTTACAGGGGTTACAGGGGTTAC<br>AGGGGTTACAGGGGTTACAGGGGTTACA-CCCCCT |
| DA260 (dsDNA) | 260 | TTACAGGGGTTACAGGGGTTACAGGGGTTACAGGGGTTACAGGGG<br>TTACAGGGGTTACAGGGGTTACAGGGGTTACAGGGGTTACAGGGG<br>TTACAGGGGTTACAGGGGTTACAGGGGTTACAGGGGTTACAGGGG<br>TTACAGGGGTTACAGGGATATCGGTTACAGGGGTTACAGGGGTTAC<br>AGGGGTTACAGGGGTTACAGGGGTTACAGGGGTTACAGGGGTTAC<br>AGGGGTTACAGGGTTACGGTTACGGTTACAGGGT |

#### Supplemental Table S2: sgRNAs used in this study

| sgRNA | sgRNA target sequence (5'-3') | PAM (5'-3') | On-target score* (%) |
|---|---|---|---|
| sgRNA1 | CATTACCCTGTTATCCC-TAC | CGG | 59.5 |
| sgRNA3 | ATTACCCTGTTATCCCT-ACC | TGG | 53.5 |
| sgRNA6 | TTACCCTGTTATCCCTA-CCT | GGG | 59.5 |
| sgRNA7 | ATTACCCTGTTATCCCT-ACC | GGG | 62.1 |
| sgRNA9 | GACATTACCCTGTTATC-CGG | GGG | 69.5 |
| sgRNA10 | CGACATTACCCTGTTAT-CCG | GGG | 61 |
| sgRNA11 (**DF254**) | ATTACCCTGTTATCCCT-AGG | GGG | 71.7 |
| sgRNA12 | CATTACCCTGTTATCCC-TAG | GGG | 64.2 |
| sgRNA13 | ACATTACCCTGTTATCC-CTA | GGG | 54.7 |
| sgRNA15 (**DA260**) | GGTTACGGTTACAGGGT-ACG | GGG | 65.7 |

*: On-target scores of sgRNAs were calculated using the sgRNA design software at the benchling.com website.

#### Supplemental Table S3: Markers used in DF254 series strain

| | Length | Sequence (5'-3') |
|---|---|---|
| Denaturing PAGE (ssDNA) | TG-81 | ATCGGTTACAGGGGTTACAGGGGTTACAGGGGTTACAGGGGTTACAGG<br>GGTTACAGGGGTTACAGGGGTTACAGGGGTTAC |
| | TG-86 | ATCGGTTACAGGGGTTACAGGGGTTACAGGGGTTACAGGGGTTACAGG<br>GGTTACAGGGGTTACAGGGGTTACAGGGGTTACAGGGG |
| | TG-91 | ATCGGTTACAGGGGTTACAGGGGTTACAGGGGTTACAGGGGTTACAGG<br>GGTTACAGGGGTTACAGGGGTTACAGGGGTTACAGGGGTTACA |
| | TG-96 | ATCGGTTACAGGGGTTACAGGGGTTACAGGGGTTACAGGGGTTACAGG<br>GGTTACAGGGGTTACAGGGGTTACAGGGGTTACAGGGGTTACAGGGGT |
| | TG-106 | ATCGGTTACAGGGGTTACAGGGGTTACAGGGGTTACAGGGGTTACAGG<br>GGTTACAGGGGTTACAGGGGTTACAGGGGTTACAGGGGTTACAGGGGT<br>TACACCCCCT |
| | CA-81 | GTAACCCCTGTAACCCCTGTAACCCCTGTAACCCCTGTAACCCCTGTAA<br>CCCCTGTAACCCCTGTAACCCCTGTAACCGAT |
| | CA-86 | CCCCTGTAACCCCTGTAACCCCTGTAACCCCTGTAACCCCTGTAACCCC<br>TGTAACCCCTGTAACCCCTGTAACCCCTGTAACCGAT |
| | CA-91 | TGTAACCCCTGTAACCCCTGTAACCCCTGTAACCCCTGTAACCCCTGTA<br>ACCCCTGTAACCCCTGTAACCCCTGTAACCCCTGTAACCGAT |
| | CA-96 | ACCCCTGTAACCCCTGTAACCCCTGTAACCCCTGTAACCCCTGTAACCC<br>CTGTAACCCCTGTAACCCCTGTAACCCCTGTAACCCCTGTAACCGAT |
| | CA-106 | AGGGGGTGTAACCCCTGTAACCCCTGTAACCCCTGTAACCCCTGTAACC<br>CCTGTAACCCCTGTAACCCCTGTAACCCCTGTAACCCCTGTAACCCCTG<br>TAACCGAT |
| Native PAGE (dsDNA) | 81 | ATCGGTTACAGGGGTTACAGGGGTTACAGGGGTTACAGGGGTTACAGG<br>GGTTACAGGGGTTACAGGGGTTACAGGGGTTAC |
| | 86 | ATCGGTTACAGGGGTTACAGGGGTTACAGGGGTTACAGGGGTTACAGG<br>GGTTACAGGGGTTACAGGGGTTACAGGGGTTACAGGGG |
| | 91 | ATCGGTTACAGGGGTTACAGGGGTTACAGGGGTTACAGGGGTTACAGG<br>GGTTACAGGGGTTACAGGGGTTACAGGGGTTACAGGGGTTACA |
| | 96 | ATCGGTTACAGGGGTTACAGGGGTTACAGGGGTTACAGGGGTTACAGG<br>GGTTACAGGGGTTACAGGGGTTACAGGGGTTACAGGGGTTACAGGGGT |
| | 106 | ATCGGTTACAGGGGTTACAGGGGTTACAGGGGTTACAGGGGTTACAGG<br>GGTTACAGGGGTTACAGGGGTTACAGGGGTTACAGGGGTTACAGGGGT<br>TACACCCCCT |

#### Supplemental Table S4: Markers used in DA260 series strain

| | Length | Sequence (5'-3') |
|---|---|---|
| Oligonucleotides (ssDNA) | TG-81 | ATCGGTTACAGGGGTTACAGGGGTTACAGGGGTTACAGGGGTTACAGG<br>GGTTACAGGGGTTACAGGGGTTACAGGGGTTAC |
| | TG-86 | ATCGGTTACAGGGGTTACAGGGGTTACAGGGGTTACAGGGGTTACAGG<br>GGTTACAGGGGTTACAGGGGTTACAGGGGTTACAGGGG |
| | TG-91 | ATCGGTTACAGGGGTTACAGGGGTTACAGGGGTTACAGGGGTTACAGG<br>GGTTACAGGGGTTACAGGGGTTACAGGGGTTACAGGGTTACGG |
| | TG-96 | ATCGGTTACAGGGGTTACAGGGGTTACAGGGGTTACAGGGGTTACAGG<br>GGTTACAGGGGTTACAGGGGTTACAGGGGTTACAGGGTTACGGTTACG |
| | TG-106 | ATCGGTTACAGGGGTTACAGGGGTTACAGGGGTTACAGGGGTTACAGG<br>GGTTACAGGGGTTACAGGGGTTACAGGGGTTACAGGGTTACGGTTACG<br>GTTACAGGGT |
| | CA-81 | GTAACCCCTGTAACCCCTGTAACCCCTGTAACCCCTGTAACCCCTGTAA<br>CCCCTGTAACCCCTGTAACCCCTGTAACCGAT |
| | CA-86 | CCCCTGTAACCCCTGTAACCCCTGTAACCCCTGTAACCCCTGTAACCCC<br>TGTAACCCCTGTAACCCCTGTAACCCCTGTAACCGAT |
| | CA-91 | CCGTAACCCTGTAACCCCTGTAACCCCTGTAACCCCTGTAACCCCTGTA<br>ACCCCTGTAACCCCTGTAACCCCTGTAACCCCTGTAACCGAT |
| | CA-96 | CGTAACCGTAACCCTGTAACCCCTGTAACCCCTGTAACCCCTGTAACCC<br>CTGTAACCCCTGTAACCCCTGTAACCCCTGTAACCCCTGTAACCGAT |
| | CA-106 | ACCCTGTAACCGTAACCGTAACCCTGTAACCCCTGTAACCCCTGTAACC<br>CCTGTAACCCCTGTAACCCCTGTAACCCCTGTAACCCCTGTAACCCCTG<br>TAACCGAT |
| Native PAGE (dsDNA) | 81 | ATCGGTTACAGGGGTTACAGGGGTTACAGGGGTTACAGGGGTTACAGG<br>GGTTACAGGGGTTACAGGGGTTACAGGGGTTAC |
| | 86 | ATCGGTTACAGGGGTTACAGGGGTTACAGGGGTTACAGGGGTTACAGG<br>GGTTACAGGGGTTACAGGGGTTACAGGGGTTACAGGGG |
| | 91 | ATCGGTTACAGGGGTTACAGGGGTTACAGGGGTTACAGGGGTTACAGG<br>GGTTACAGGGGTTACAGGGGTTACAGGGGTTACAGGGTTACGG |
| | 96 | ATCGGTTACAGGGGTTACAGGGGTTACAGGGGTTACAGGGGTTACAGG<br>GGTTACAGGGGTTACAGGGGTTACAGGGGTTACAGGGTTACGGTTACG |
| | 106 | ATCGGTTACAGGGGTTACAGGGGTTACAGGGGTTACAGGGGTTACAGG<br>GGTTACAGGGGTTACAGGGGTTACAGGGGTTACAGGGTTACGGTTACG<br>GTTACAGGGT |

**Supplemental Table S5: Oligonucleotide markers (OM) used in native PAGE**

| Oligos | Sequence (5’-3’) |
|---|---|
| 466 106 0nt F | ATCGGTTACAGGGGTTACAGGGGTTACAGGGGTTACAGGGGTTACAGGGG<br>TTACAGGGGTTACAGGGGTTACAGGGGTTACTCGGGTTACACGGGTTACAC<br>CCCCT |
| 467 106 0nt R | AGGGGGTGTAACCCGTGTAACCCGAGTAACCCCTGTAACCCCTGTAACCCC<br>TGTAACCCCTGTAACCCCTGTAACCCCTGTAACCCCTGTAACCCCTGTAACC<br>GAT |
| 468 106 10nt F | ATCGGTTACAGGGGTTACAGGGGTTACAGGGGTTACAGGGGTTACAGGGG<br>TTACAGGGGTTACAGGGGTTACTCGGGTTACACGGGTTACAGGGGTTACAC<br>CCCCT |
| 469 106 10nt R | ACCCCTGTAACCCGTGTAACCCGAGTAACCCCTGTAACCCCTGTAACCCCT<br>GTAACCCCTGTAACCCCTGTAACCCCTGTAACCCCTGTAACCGAT |
